# Proteogenomics analysis of non-coding region encoded peptides in normal tissues and five cancer types

**DOI:** 10.1101/2020.04.10.029306

**Authors:** Rong Xiang, Leyao Ma, Mingyu Yang, Zetian Zheng, Xiaofang Chen, Fujian Jia, Fanfan Xie, Fuqiang Li, Kui Wu, Yafeng Zhu

## Abstract

Previous proteogenomics studies have identified peptides encoded by non-coding sequences such as pseudogenes and long non-coding RNAs (lncRNAs) in healthy human tissues as well as in cancers. However, these studies are either limited to analyze only healthy or cancerous tissues, lacking direct comparison between them. In this study, we used an established proteogenomics analysis workflow to analyze proteomics data from 926 cancer samples of five cancer types and 31 different healthy human tissues. We observed the protein level expression of pseudogenes can be classified as ubiquitous or lineage expression. The ubiquitously translated pseudogenes are homologous to house-keeping genes. Our results suggest there is common mechanism underlying the translation of pseudogenes in both normal and tumors. Moreover, we discovered several translated non-coding genes such as *DGCR5* and *RHOXF1P3* that were up-regulated in tumors compared to normal. These translated pseudogenes imply the biological function of pseudogenes extends to protein level yet to be studied. Further, from the non-coding region encoded peptides specifically detected in tumors we have predicted a large number of potential neoantigens which can be developed as cancer vaccine.

## Introduction

Recently, many mass spectrometry based proteomics studies have identified peptides from non-coding regions of human genome^1–5^. Some peptides are identified from genomic regions in proximity to protein-coding genes, which indicate incorrect exon boundary or a missed exon. Others are identified from currently annotated non-coding sequences include pseudogenes, lncRNAs, protein-coding gene’s untranslated region, alternative reading frame or anti-sense strand.

It is conventionally believed that pseudogenes have lost protein-coding functions due to accumulated deleterious mutations. Recently, the analysis of RNA-seq data from cancer cell lines and tumors has showed active transcription of pseudogene in different cell lineages and cancer types, and some pseudogenes have shown cancer specific expression compared to normal tissues^6–8^. In addition to RNA level detection of pseudogene expression, several independent proteomics studies have identified peptide evidence of pseudogenes and lncRNAs translation in normal tissues and cancer cell lines^1–3^. It remains unknown if pseudogenes are translated in tumor tissues and whether pseudogene translation is a sporadic event or under certain specific regulation in different type of tumors.

Here, we analyzed publicly available proteomics data from 926 cancer samples of five cancer types, and 31 different healthy human tissues using previously developed proteogenomics pipeline^4^. With the data, we aim to find out what non-coding sequences including pseudogenes/lncRNAs are actively translated in healthy and tumor tissues and if they exert tissue specific or cancer specific expression. Secondly, a published study by Laumont et al detected more tumor specific antigens from non-coding regions compared to mutations in protein-coding regions^9^. The *in vivo* mice experiments demonstrated immunization against these non-coding region peptides could prevent tumors in mice that are received oncogenic cancer cells. Inspired by this, our second goal is to investigate whether these pseudogene/lncRNA encoded proteins can be classified as tumor specific antigens which can potentially utilized as cancer vaccines.

## Results

### Proteomics detects ubiquitous and tissue-specific translation of pseudogenes

We downloaded proteomics data of 40 normal samples from 31 healthy tissues and 926 cancer samples from 5 cancer types from the PRIDE database and CPTAC Data Portal. The websites where the datasets are downloaded are listed in supplementary table S1. To search the proteomics data, we first constructed a core database including ENSEMBL human proteins, and peptide sequences from three frame translation of annotated pseudogenes from GENCODE v28 and lncRNA from LNCpedia 4.1. This core database was used as search database for proteomics data of healthy tissue. As for different cancer datasets, cancer specific mutant proteins’ sequences are supplemented to the core search database (See details in Method). The proteogenomics search was performed using a previously published pipeline^4^ (Figure S1).

In total, we identified 7882 novel peptides from 31 normal tissues and 9013 novel peptides from five cancer types at 1% class specific FDR. Novel peptides are defined as peptide sequences that are absent in current known protein databases (Uniprot human reference proteome plus GENCODE v28 human protein database). The number of unique peptides per novel coding loci (peptides within 10 kb distance were grouped into one locus) were summarized for 31 healthy tissues, and 13 tumor datasets respectively (Figure 1a and 1b). The novel loci were divided into three groups according to whether they are supported by one, two and more than two unique peptides. Removing loci supported by single peptide, 687 novel coding loci was identified of which 131 overlap with the 220 identified from normal tissues (Figure 1a).

**Figure 1.**
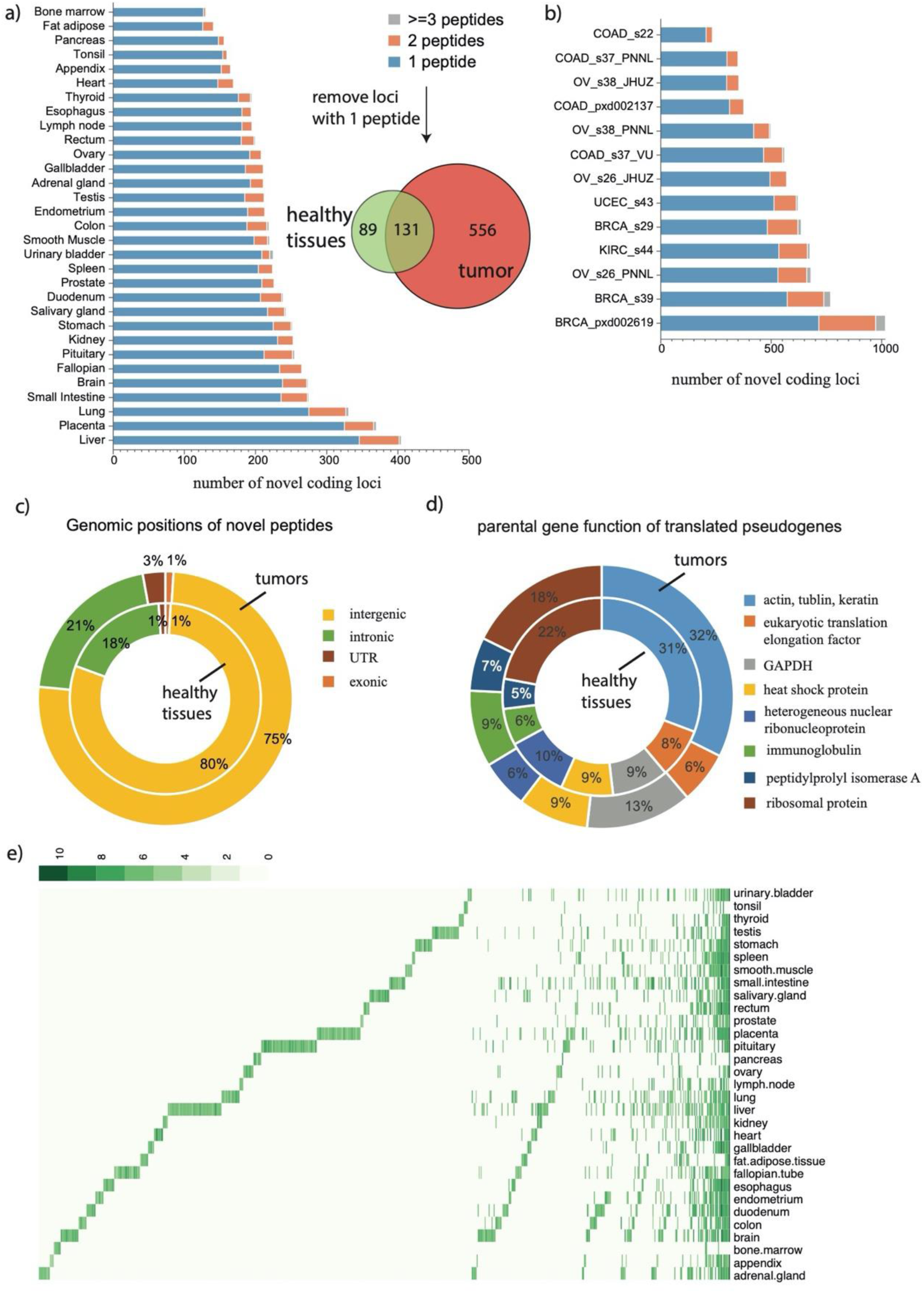
Non-coding gene encoded peptides detected from normal tissues and tumor datasets. a) the number of novel coding loci detected in 31 normal tissues (peptides are grouped to one locus if they are encoded by the same non-coding gene or if they locate within 10 kb distance). Venn diagram shows the overlap of novel coding loci (>= 2 unique peptides detected) between normal tissues and tumors. b) the number of novel coding loci detected in 13 cancer datasets. c) annotation of genomic positions where non-coding gene encoded peptides are detected. d) annotation of parental genes’ function of translated pseudogenes. e) heatmap of MS1 intensity of non-coding gene encoded peptides detected in normal tissues.

Next, we categorized the novel-coding loci based their genomic positions (Figure 1c). This result indicates tumor and normal has no major differences of expressed novel coding loci in terms of genomic regions. Our further analysis discovered the majority of novel coding loci in intergenic and intronic regions are derived from pseudogenes. We therefore annotated the parental genes’ functions of translated pseudogenes (Figure 1d). In consistent to the findings revealed by previous RNA-seq data analysis^6^, the frequently detected pseudogenes in healthy tissues and tumors are homologous to house-keeping genes such as cytoskeleton proteins (actin, keratin, tubulin), ribosomal proteins, nuclear ribonucleoproteins, heat shock proteins and eukaryotic translation elongation factor, peptidylprolyl isomerase (Figure 1d, supplementary figure S2 a). Our results showed that there is a common mechanism that regulates the translation of pseudogenes in healthy and tumor tissues.

Previous proteogenomic studies have rarely performed quantitative analysis, here we quantified non-coding region encoded peptides by MS1 maximum peak intensity using moFF^10^. We limited the analysis to novel coding loci with at least two unique peptides. Our analysis identified three groups of pseudogene expression: ubiquitous, non-specific and tissue specific expression (Figure 1e). The ubiquitously translated pseudogenes are those homologous to house-keeping genes mentioned above. Most pseudogenes exhibited non-specific expression, which are robustly translated in one or two tissues but frequently translated at lower levels in other tissues. This is similar to the previously reported transcription level expression pattern of pseudogenes^6^. A few representative tissue-specific pseudogenes/ lncRNAs are listed in Table 1. We detected two previously reported tissue specific non-coding gene translation, testis specific *TATDN2P1* (TatD DNase domain containing 2 pseudogene 1, supported by two unique peptides) and placenta specific lncRNA *lnc-CACNG8-28:1* (supported by eight unique peptides)^4^. In addition, some more novel tissue specific non-coding gene encoded peptides were discovered in different tissues (supplementary table S2). For example, ten unique peptides encoded by a lncRNA *lnc-AFF3-13:1* located in 5’ UTR of gene *TSGA10* were detected in fallopian. Six unique peptides from *PRH1-PRR4* read-through transcript were detected in salivary gland. Pseudogene *CCDC150P1* were detected with five unique peptides in testis, and this pseudogene *CCDC150P1* transcript is also specifically expressed in testis according to GTex data (supplementary figure S2 b). Interestingly, in pituitary tissue, peptides were identified from lncRNAs that overlap with the coding exons of two pituitary specific protein coding genes, *GH1* and *POMC*, but in non-canonical reading frames. Our data indicates the two genes may have dual coding frames that encode unknown new proteins.

**Table 1.** translated pseudogene/lncRNA showed tissue specific expression

| ncRNA ID | Description | Detected in tissue | # detected peptides |
| --- | --- | --- | --- |
| lnc-AFF3-13:1 | Located in 5' UTR of gene TSGA10 | fallopian | 10 |
| lnc-CACNG8-28:1 | lncRNA ENSG00000267943, CTD-2620I22.3 | placenta | 8 |
| lnc-M6PR-48:14 | PRH1-PRR4 read-through transcript | salivary gland | 6 |
| CCDC150P1 | Coiled-Coil Domain Containing 150 Pseudogene 1 | testis | 5 |
| AKR1B1P7 | aldo-keto reductase family 1 member B1 pseudogene 7 | adrenal gland | 3 |
| CBX1P1 | chromobox 1 pseudogene 1 | fallopian | 3 |
| ANXA2P3 | Annexin A2 Pseudogene 3 | placenta | 3 |
| RP6-218J18.2 | processed pseudogene, similar to RNA binding motif protein 39 | adrenal gland | 2 |
| PFN1P2 | profilin 1 pseudogene 2 | appendix | 2 |
| UBBP4 | ubiquitin B pseudogene 4 | fallopian | 2 |
| AKR1C8P | aldo-keto reductase family 1 member C8, pseudogene | liver | 2 |
| CES1P2 | carboxylesterase 1 pseudogene 2 | liver | 2 |
| lnc-CABIN1-20:1 | Novel transcript, AL928654.3 | lung | 2 |
| DDX18P6 | DEAD-box helicase 18 pseudogene 6 | ovary | 2 |
| IMMTP1 | inner membrane mitochondrial protein pseudogene 1 | pituitary | 2 |
| AC138932.1 | polycystic kidney disease 1 (PKD1) pseudogene | pituitary | 2 |
| lnc-CYB561-18:1 | non-canonical reading frame of GH1 (Growth Hormone 1) | pituitary | 3 |
| lnc-DNAJC27-5:1 | non-canonical reading frame of POMC | pituitary | 2 |
| lnc-MYOG-56:3 | non-canonical reading frame of TNNT2 | heart | 2 |
| HSD17B1P1 | Hydroxysteroid 17-Beta Dehydrogenase 1 Pseudogene 1 | placenta | 2 |
| TATDN2P1 | TatD DNase domain containing 2 pseudogene 1 | testis | 2 |
| lnc-C8orf86-8:1 | Intergenic, C8orf86(dist=12750), RNF5P1(dist=58763) | gallbladder | 2 |

### Recurrent detection of non-coding gene encoded peptides in different cancers

As a way to evaluate the reproducibility of the detected non-coding gene encoded peptides, we analyzed the overlap of detected novel coding loci in different datasets within and between cancer types (Figure 2). We found the most novel coding loci with two-unique peptides supported in breast cancer. Among the downloaded datasets, CPTAC studies generated more pseudogene identifications, likely a result of high-quality data and larger cohort size. Among different cancer types, the repeatedly detected pseudogenes are those homologous to house-keeping genes including pseudogenes of eukaryotic elongation factors, GAPDH, actin/keratin/tubulin, heat shock protein family, heterogeneous nuclear ribonucleoprotein and ribosomal proteins. These pseudogenes are also ubiquitously expressed in different healthy tissues.

**Figure 2.**
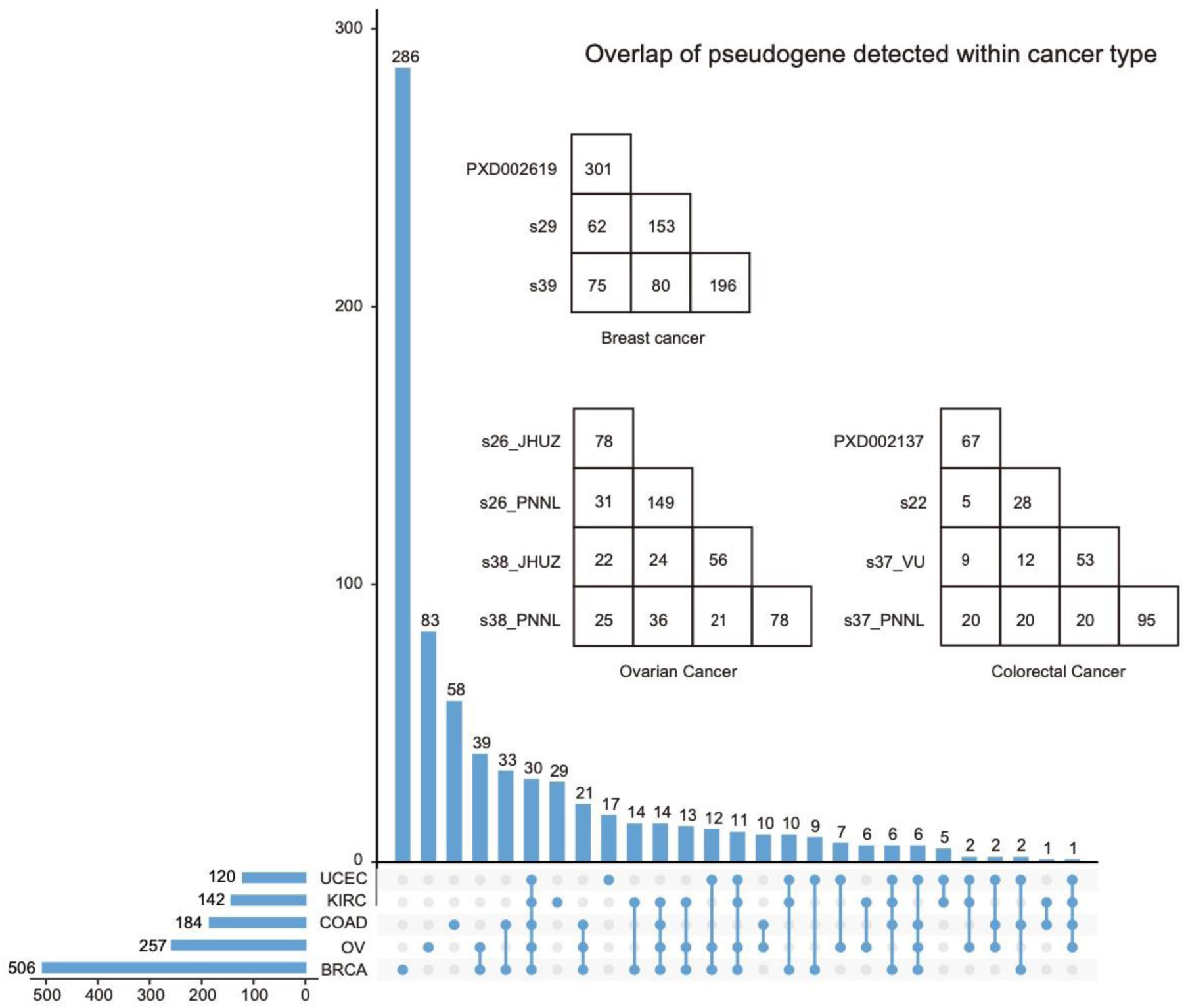
Overlap of novel coding loci detected in different cancers. The horizontal bars indicate the total number of novel coding loci (>= 2 unique peptides) detected in each cancer combining all datasets of this cancer type. Vertical bars with only one dot below the axis indicate the number of novel loci uniquely identified in the corresponding cancer types. Vertical bars with multiple dots connected below the axis indicate the number of novel loci detected in all corresponding cancer types, and absent in remaining cancer types. For example, the vertical bar with the number 286 on it indicates 286 novel loci uniquely found in BRCA datasets. The vertical bar with the number 39 means 39 of the novel loci commonly detected in BRCA and OV datasets are absent in other cancer types. The vertical bar with the number 30 means 30 novel loci commonly detected in all five cancer types. The last vertical bar with the number 1 means only one of the novel loci commonly detected in OV, COAD, KIRC and UCEC is absent in BRCA datasets. In the matrix, the number in the last column of each row indicate the number of novel coding loci identified in individual datasets, the others show the overlap of two corresponding datasets.

Next, we looked the overlap of novel coding loci detected within same cancer type. Apart from the ubiquitously expressed pseudogenes, many pseudogenes were recurrently detected in specific cancers (Table 2). The notable examples are *RHOXF1P3, PA2G4P4* and *MCTS2P* which are repeatedly detected from independent datasets of breast and ovarian cancers and supported by multiple unique peptides.

**Table 2.** translated pseudogenes detected in cancer

| Pseudogene | Description | Detected in cancer(s) | # detected peptides |
| --- | --- | --- | --- |
| *RHOXF1P3* | Rhox homeobox family member 1 pseudogene 3 | BRCA, OV | 8 |
| *PA2G4P4* | proliferation-associated 2G4 pseudogene 4 | BRCA, CCRCC | 6 |
| *MCTS2P* | malignant T cell amplified sequence 2 pseudogene | BRCA, OV, UCEC | 4 |
| *GLYATL1P4* | glycine-N-acyltransferase like 1 pseudogene 4 | BRCA | 3 |
| *NPM1P52* | nucleophosmin 1 pseudogene 52 | BRCA | 3 |
| *AC051619.2* | H3 histone family 3A H3F3A pseudogene | OV | 2 |
| *AL158201.1* | phosphoglycerate kinase 1 PGK1 pseudogene | OV, CCRCC | 4 |
| *AC104297.1* | pyrophosphatase inorganic 1 PPA1 pseudogene | OV | 2 |
| *VDAC1P7* | voltage dependent anion channel 1 pseudogene 7 | OV | 2 |
| *AC013470.3* | Sjogren syndrome antigen B (SSB) pseudogene | OV | 3 |
| *PHBP9* | prohibitin pseudogene 9 | OV | 2 |
| *NAP1L4P3* | nucleosome assembly protein 1 like 4 pseudogene 3 | OV | 2 |
| *AL109618.1* | NS1-associated protein 1 (NSAP1) pseudogene | OV | 2 |
| *STIP1P3* | stress induced phosphoprotein 1 pseudogene 3 | OV | 2 |
| *ILF2P1* | interleukin enhancer binding factor 2 pseudogene 1 | COAD | 2 |
| *AC018797.3* | WD repeat domain 12 (WDR12) pseudogene | COAD | 2 |
| *FTH1P2* | ferritin heavy chain 1 pseudogene 2 | COAD | 2 |
| *AL359263.1* | adaptor-related protein complex 2, beta 1 subunit (AP2B1) pseudogene | COAD | 2 |
| *AC083873.1* | suppression of tumorigenicity 13 (ST13) (Hsp70 interacting protein) pseudogene | UCEC, CCRCC, BRCA | 3 |
| *AC231533.2* | acetyl-Coenzyme A acyltransferase 2 (ACAA2) pseudogene | CCRCC | 4 |
| *AL583856.1* | spermine synthase (SMS) pseudogene | CCRCC | 2 |
| *GM2AP1* | GM2 ganglioside activator pseudogene 1 | CCRCC | 2 |

In addition to pseudogenes, we also found several long non-coding RNA encoded peptides were detected in specific cancers (Table 3). For example, the non-coding RNA genes *DGCR5* (DiGeorge Syndrome Critical Region Gene 5, located on chromosome 22q11), also known as LINC00037, were detected with ten unique peptides in clear cell renal cell carcinoma (CCRCC). Previous study by Gong et al reported *DGCR5* is ubiquitously expressed in normal tissues, with much higher expression in CCRCC compared to normal kidney tissues^11^. Our analysis showed that *DGCR5* probably encodes a functional protein product in CCRCC.

**Table 3.** translated lncRNA detected in cancer

| lncRNA | Annotation | Detected in cancer(s) | # detected peptides |
| --- | --- | --- | --- |
| DGCR5 | DiGeorge Syndrome Critical Region Gene 5 (Non-Protein Coding), also known as LINC00037, lnc-SERPIND1-41:10 | CCRCC | 10 |
| lnc-MMADHC-67:1 | KCNJ3 upstream (distance=532 bp) | UCEC | 2 |
| LINC02043 | lnc-EPAS1-55:1 (lncpedia ID) | BRCA, | 2 |
| OSMR-AS1 | OSMR Antisense RNA 1 | Colon cancer | 2 |
| lnc-ARMT1-2:1 | in the intron of RMND1 | BRCA | 3 |
| LINC02210 | lnc-CCDC103-52:14 (lncpedia ID) | BRCA | 3 |
| lnc-GLRX5-72:1 | in the intron of TRIP11 | BRCA | 4 |
| HCG4B | HLA Complex Group 4B (Non-Protein Coding) | BRCA | 2 |

Since pseudogene expression has been extensively analyzed at transcript level using RNA-seq data^6,7^ and the major biological functions of pseudogenes are revealed at RNA level, we wonder what pseudogenes expressed at RNA level are translated into proteins. Therefore, we compared pseudogenes detected in our proteomics analysis with two major studies in which expression of pseudogenes were investigated through RNA-seq analysis^6,7^. The pseudogenes detected both in RNA and protein level are pseudogenes of house-keeping genes such as ribosomal proteins, GAPDH, cytokeratin, eukaryotic translation initiation factors and heterogeneous nuclear ribonucleoprotein. In addition, pseudogenes corresponding to cancer associated genes *HMGB1*, *VDAC1* and *PTMA* reported in previous RNA-seq study^6^ were detected both in healthy tissues and cancers in our proteomics analysis. In comparison, many of the known functional pseudogenes such as *PTENP1* were not detected in proteomics data, this is not a surprise since they are functional as ceRNA molecules regulating the expression of their parental genes^12^. Another example is the breast cancer pseudogene *ATP8A2P1* which showed high expression at RNA level^6,7^ was not detected at protein level in any of the breast cancer proteomics data, suggesting this pseudogene probably only exert functions at RNA level.

### Differential expressed non-coding gene encoded peptides between tumor and normal

We investigated if certain pseudogenes/lncRNA peptides have elevated abundance in tumors in a COAD dataset with 8 paired colon cancer samples and matched normal tissues (PXD002137)^13^. In this data, 73 pseudogenes/lncRNA were identified supported by multiple peptides. We removed peptides identified in only one sample, leaving 71 pseudogenes/lncRNA. Samples with missing values were imputed using left-shifted Gaussian distribution. A heatmap of 25 significant pseudogenes/lncRNA (p. adj<0.05, Limma) with the data centered per protein is shown (Figure3a, supplementary table S3). Most of significant pseudogenes/lncRNA are upregulated. For example, *lnc-KMT5B-20:1*, *MKKS* 5’UTR and *ACTBP2* are up-regulated in colon cancer (Figure 3b).

**Figure 3.**
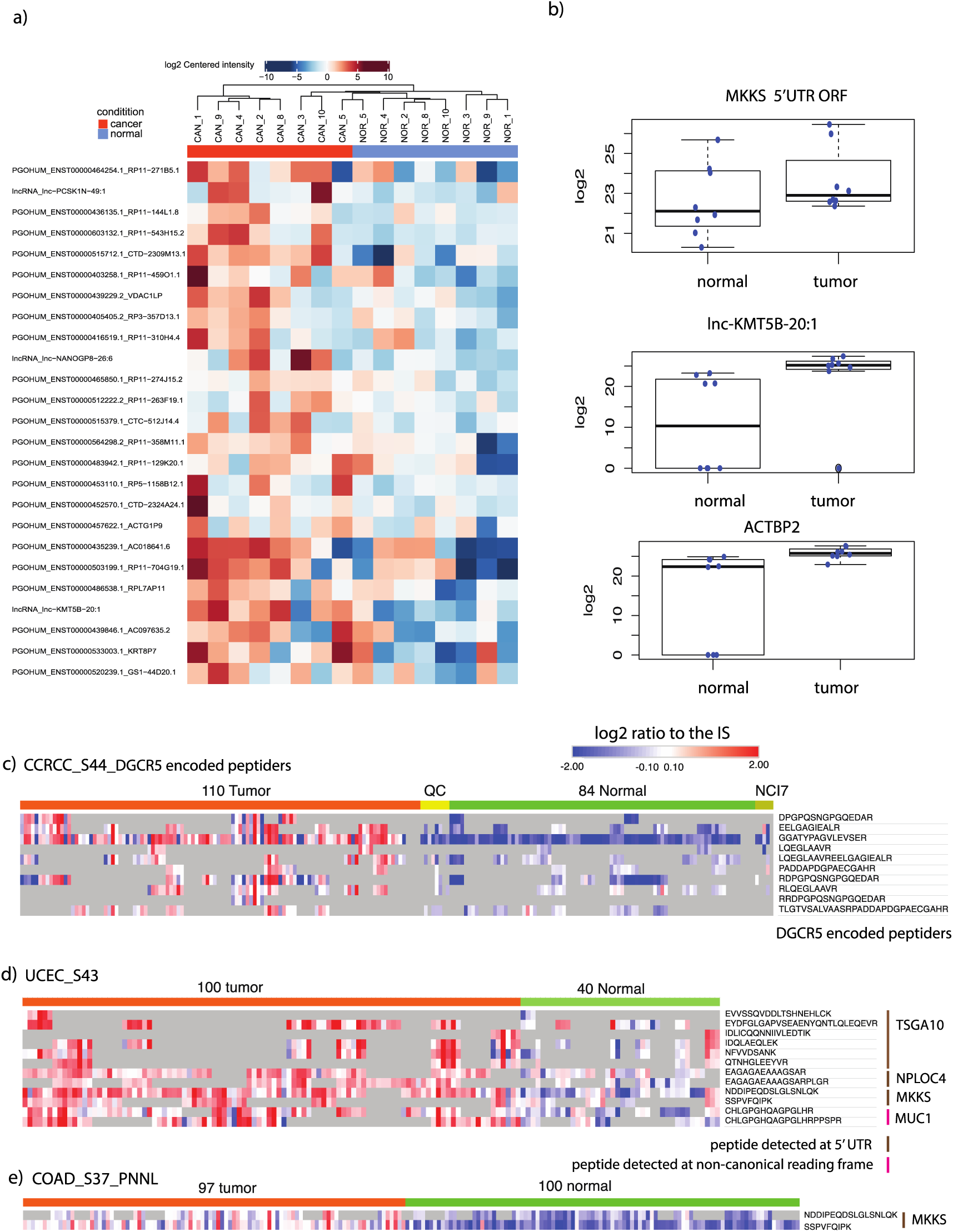
Non-coding region encoded peptides detected in tumor. a) heatmap of log2 centered intensity of peptides detected from non-coding genes in COAD (PXD002137). b) boxplot of non-coding genes significantly differentially expressed between tumor and normal (p.adj<0.05). c) relative expression of the ten unique peptides detected from DGCR5 in tumor (CCRCC) and normal. d) relative expression of peptides detected at 5’ UTR in tumor (UCEC) and normal. e) relative expression of peptides detected from MKKS 5’ UTR in tumor (COAD) and normal.

In other cancer datasets, we also detected several non-coding gene encoded peptides show increased expression in tumors. For examples, the peptides encoded by non-coding gene *DGCR5* showed significantly higher expression level in CCRCC compared to adjacent normal tissues (Figure 3c). This contradicts the finding at RNA level where *DGCR5* is ubiquitously expressed in normal tissues^11^. In Uterine Corpus Endometrial Carcinoma (UCEC), peptides detected from 5’ UTR or non-canonical reading frame of protein coding genes, *TSGA10*, *NPLOC4*, *MKKS* and *MUC1* were more abundant in tumors compared to normal tissues (Figure 3d). Similarly, increased expression of peptides from *MKKS* 5’ UTR was also detected in another colon adenocarcinoma (COAD) dataset (Figure 3e).

### Pseudogene RHOXF1P3 expression is associated with ER positive breast tumors

Next, we tried to correlate the expression of non-coding gene encoded peptides with tumor subtypes. Interestingly, the pseudogene *RHOXF1P3* detected with eight unique peptides showed increased expression (2 to 16 fold up-regulation) in a subset of estrogen receptor (ER) positive breast cancer from CPTAC breast cancer Discovery cohort (Figure 4a-4b)^14^. *RHOXF1P3* peptides were also detected in CPTAC breast cancer confirmatory cohort, expressed in 13 out of 133 breast tumor and 2 out of 18 breast normal samples (Figure 4c). In addition, *RHOXF1P3* encoded peptides were also detected in two ovarian cancer patients (Figure 4d). We then analyzed the expression of *RHOXF1P3* in a published RNA-seq data including 63 breast tumors and 10 adjacent normal tissues (Figure 4e). These combined results indicate *RHOXF1P3* is upregulated in breast tumors and it is likely to be associated with ER signaling in breast and ovarian cancer.

**Figure 4.**
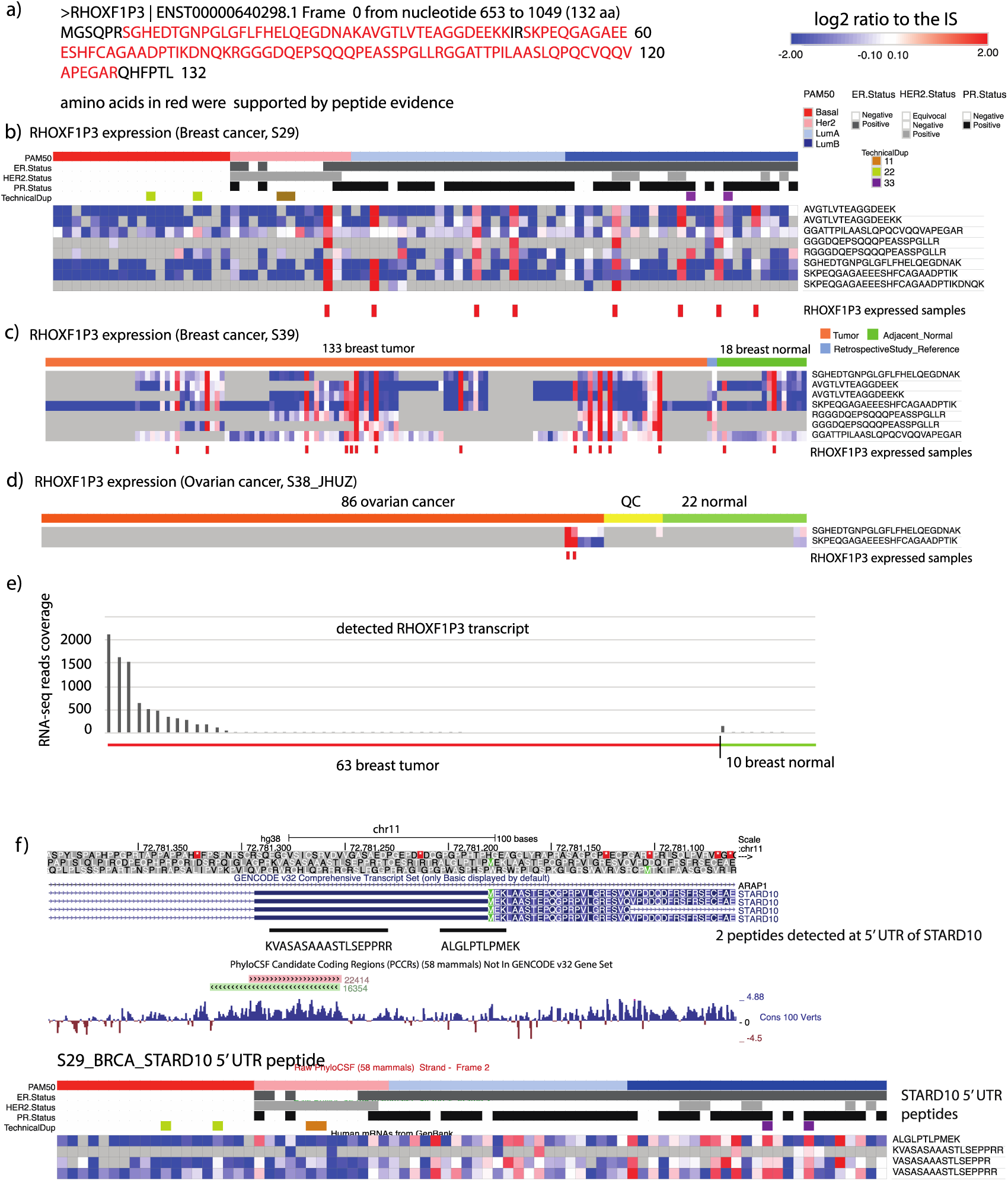
Non-coding regions encoded peptides associated with ER positive breast tumor. a) predicted protein sequence of pseudogene RHOXF1P3 (amino acids in read were supported by detected peptides). b) relative expression of RHOXF1P3 encoded peptides in 77 breast tumors. Grey color boxes indicate missing values. c) relative expression of RHOXF1P3 encoded peptides in 133 breast tumors and 18 adjacent normal tissues. d) relative expression of RHOXF1P3 encoded peptides in 86 ovarian tumors and 22 normal ovarian tissues. e) RNA seq read count of RHOXF1P3 transcript in breast tumor and normal. f) relative expression of peptides detected at 5’ UTR of STARD10 in 77 breast tumors.

Meanwhile, we found another example. Peptides detected from 5’ UTR of *STARD10* also displayed higher abundance in ER positive breast tumors (Figure 4f). *STARD10* is a lipid transfer protein and this protein has been previously reported to be overexpressed in breast cancers and correlate ErbB2/Her2 status ^15^. Our data suggests that this gene may use an upstream non-AUG start codon to initiate translation in ER positive breast tumors.

### LINE-1 retrotransposon ORF1 encoded peptides show higher expression in tumors

As evidenced in many studies, cellular mechanism that repress the expression of repetitive DNA is disrupted in cancer cells. Overexpression of satellite repeats were previously observed in pancreatic and other epithelial cancers^16,17^. This phenomenon correlates with the overexpression of the long interspersed nuclear element 1 (LINE-1) retrotransposon, which is suggested as a hallmark of many cancers^17^.

In previous proteogenomics studies, peptides mapped to multiple genomic locations were often neglected. In our analysis, LINE-1 retrotransposon ORF1 encoded peptides were detected in different tumor datasets. LINE-1 RNA contains two non-overlapping open reading frames, encoding two proteins ORF1p and ORF2p. The expression level of ORF1p is 1000-10,000 times higher than ORF2p^18^. In the analyzed proteomics data, we detected ORF1p peptides from all five cancer types, and ORF2p peptides from CCRCC (supplementary table S4). The quantitative analysis showed higher expression of the LINE-1 ORF1p encoded peptides in tumors compared to normal samples (Figure 5). In comparison, ORF2p peptides were rarely detected. It corroborates findings from an antibody-based study which concluded that LINE-1 ORF2p expression is hardly detectable in human cancers ^19^. Surprisingly, we also detected peptides of LINE-1 ORF1 in healthy tissues, including lung, ovary and prostate (supplementary table S5). This may be explained by a recent study which showed LINE-1 activity becomes derepressed in senescent cells 20 and healthy tissues could have senescent cells at old age.

**Figure 5.**
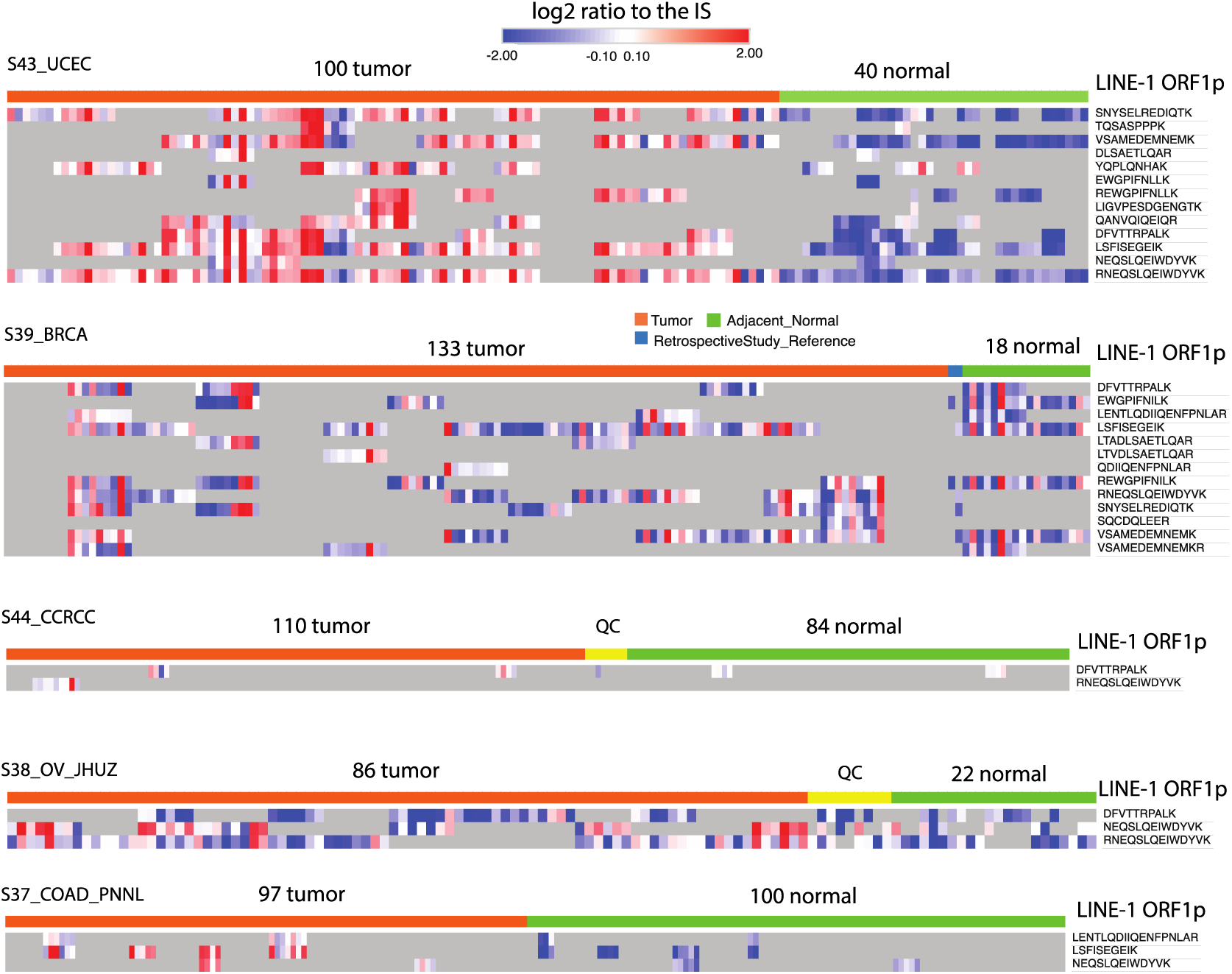
Relative abundance of LINE-1 ORF1 encoded peptides in tumor and normal. The heatmap shows the log2 relative abundance of LINE-1 ORF1 encoded peptides in five different cancers. Grey boxes indicate missing values.

### Non-coding region encoded peptides as a new class of tumor neoantigen

Laumont *et al* demonstrated non-coding region encoded peptides can be used as cancer vaccine to prevent tumor progression^9^. Here, we try to predict if any of non-coding region encoded peptides can be used as potential tumor neoantigen. The process of tumor neoantigen presentation based on the tumor specific novel peptide mainly depends on: 1) whether the novel peptide can be selectively hydrolyzed by proteasome; 2) whether the digested peptide can be selectively transported by antigen processing-related transporters;3) if the peptides can bind to MHC molecules, and can the presented antigen be recognized by T cells. Therefore, novel tumor-specific peptides were predicted by combining NetCTLpan and NetMHCpan, both intracellular presentation of MHC Ⅰ class antigens and the ability of MHC and extracellular binding of antigens are considered.

After filtering out all novel peptides expressed in healthy tissues, 1587 peptides from 524 pseudogenes were used in neoantigen prediction (table S6). By NetMHCpan and NetCTLpan 4.0 prediction, 444 and 296 pseudogenes/lncRNAs have at least one 9-mer peptide with predicted affinity ranked at threshold ≤0.5% and ≤1%, respectively (Figure 6a). Taking the overlap novel loci of the two softwares, 296 novel loci encode peptides predicted to be presented by at least one HLA allele and recognized by T cells (Figure 6b, supplementary table S7). Some examples of predicted neoantigens from non-coding genes are shown in Table 4. In GTEx-data, the transcripts that encode these antigen peptides are poorly expressed or not expressed in healthy tissues (Figure 6c), allowing them to be explored as peptide vaccine to treat cancer^9^.

**Table 4.**
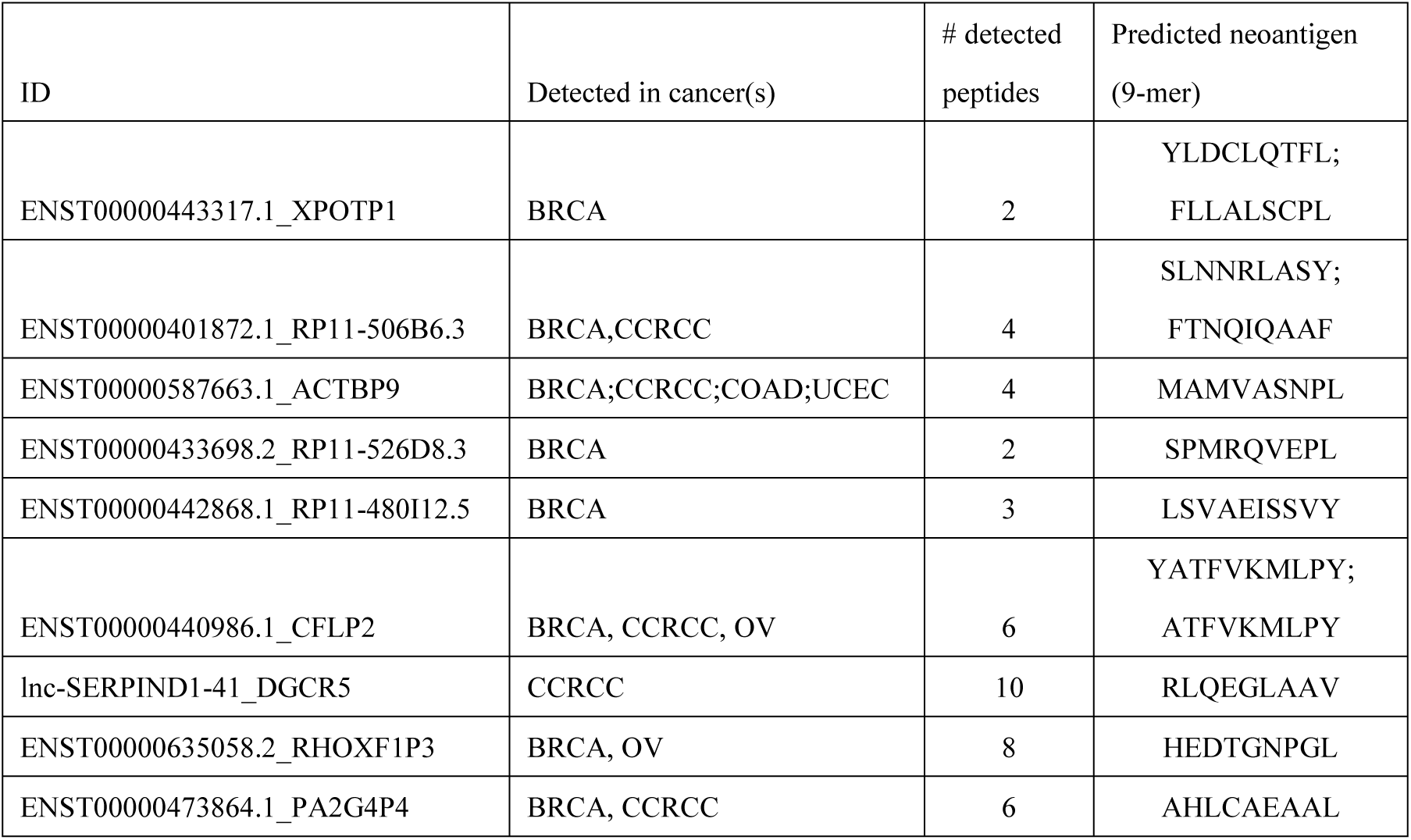
predicted neoantigen from non-coding genes

**Figure 6.**
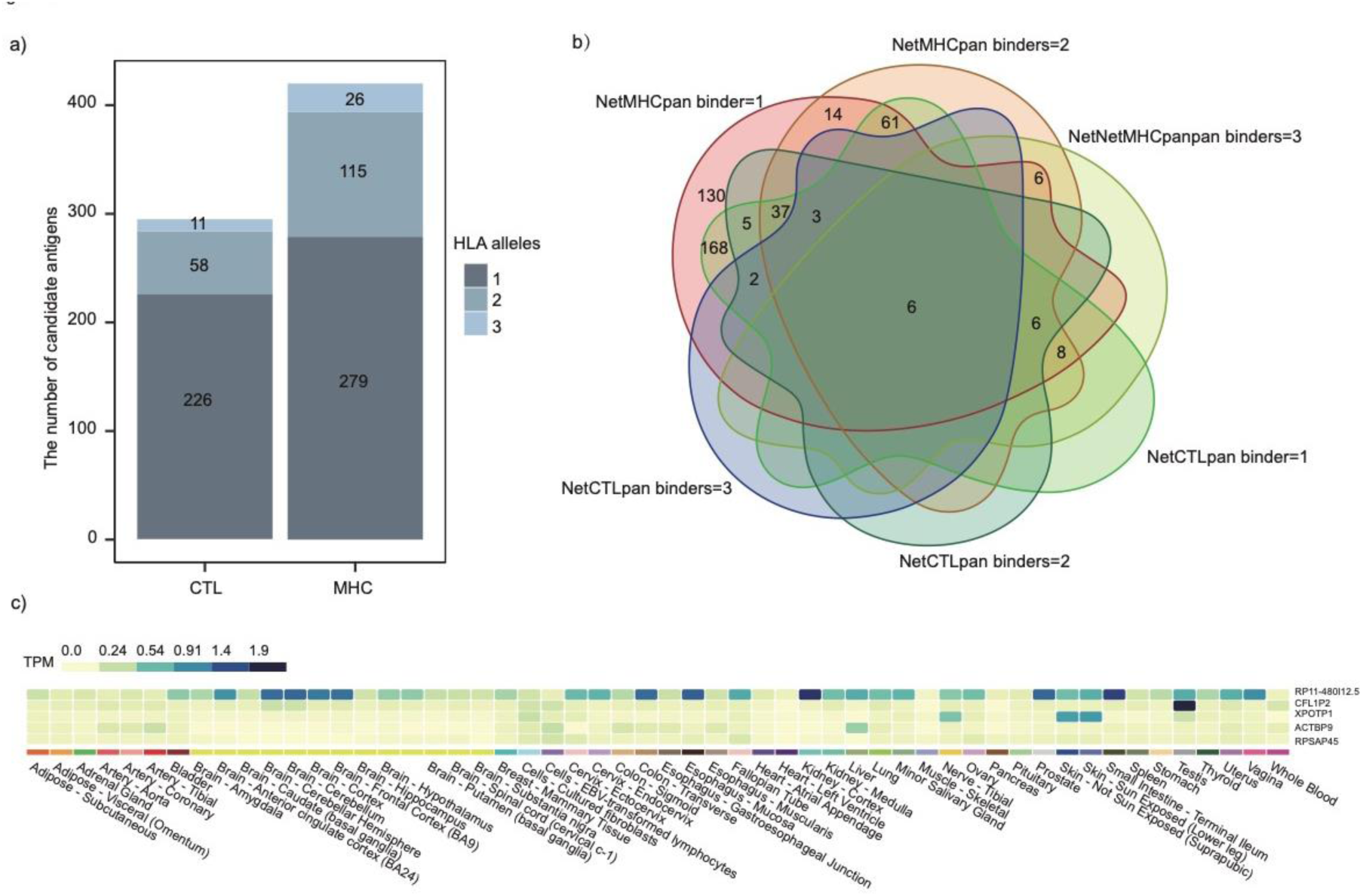
Neoantigens are derived from non-coding regions. a) distribution of the candidate novel loci predicted by NetMHCpan and NetCTLpan (classification was based on the number of binding with HLA allele(s)). b) the overlapping candidate novel loci predicted by NetMHCpan and NetCTLpan. c)RNA level expression of six candidate novel loci that encode peptides predicted to bind with at least three HLA alleles in healthy tissues (GTEx data).

## Discussion

Protein-coding genes have been dominantly annotated based on RNA-level data ^21^. Several studies integrating the ribosome profiling data to unbiasedly search all potential coding sequences led to the findings that many of the non-coding regions of human genome are being translated^22–24^. Since then, non-coding regions encoded peptides have been detected in several large scale of mass spectrometry-based proteomics studies ^4,25^. Here, we analyzed a large amount of high-quality mass spectrometry proteomics data from 31 healthy normal tissues and five cancer subtypes. In our analysis, we detected ubiquitous and tissue-specific expression of pseudogenes at the peptide levels in normal and cancer tissues, including 687 novel coding loci identified from five tumors and 220 novel loci identified from normal tissues. These novel coding loci are supported by at least two peptides and mainly derived from pseudogenes. The ubiquitously translated pseudogenes are homologous to house-keeping genes. Most pseudogenes exhibit nonspecific expression, showing strong translation in one or two tissues and low translation in other tissues. This is similar to the previously reported transcriptional expression pattern of pseudogenes.

A few tissue-specific or cancer-specific pseudogene/lncRNA were found. *CCDC150P1* is expressed specifically in the testis at both transcriptional and protein levels. Non-coding RNA *DGCR5* encoded peptides was detected in CCRCC and showed higher expression compared to normal tissues. We also detected previously reported testis specific *TATDN2P1* and placenta specific lncRNA *lnc-CACNG8-28:1*^4^. Many pseudogenes have been found repeatedly in specific cancers and supported by multiple unique peptides, such as Rhox homeobox family member 1 pseudogene 3 (*RHOXF1P3*), proliferation-associated 2G4 pseudogene 4 (*PA2G4P4*) and malignant T cell amplified sequence 2 pseudogene (*MCTS2P)*. It’s worth noting that *RHOXF1P3* detected with eight unique peptides were upregulated in a subset of estrogen receptor (ER) positive breast cancer. It suggests that the translation of *RHOXF1P3* may be associated with ER signaling in breast and ovarian cancer. According to the predicted results by the NetMHCpan4.0 and NetCTLpan, there are a large number of potential tumor specific antigens from the non-coding regions.

Our results suggest there is common underlying mechanism that regulate the translation of non-coding sequences such as pseudogenes in both normal and tumors. In addition, non-coding region encoded peptides specifically detected in tumors or in certain cancer subtypes suggest non-coding region encoded proteins/peptides also play a role in cancer and their biological functions are yet to be investigated. In the aspect of clinical applications, these peptides detected from non-coding regions, especially the ones specific to tumors, may provide a new class of tumor neoantigens possessing more potent immunogenic activity to be explored as cancer vaccine ^9^.

## Material and methods

### Data sets

LC-MS/MS raw files of 40 normal samples from 31 healthy tissues and 926 cancer samples from five cancer types were obtained from National Cancer Institute Clinical Proteomic Tumor Analysis Consortium (CPTAC) and PRoteomics IDEntifcations (PRIDE) database. See supplementary table S1 for details.

### Database construction

For the healthy tissues, the hypothetical peptide database contains the human protein database of Ensembl 92 and peptide sequences from three frame translation of annotated pseudogenes and lncRNAs. Pseudogenes were downloaded from GENCODE v28 including both annotated and predicted. LncRNAs were downloaded from LNCpedia 4.1. For each cancer type, mutations derived protein sequence variants extracted from the Cancer Genomic Data Server (CGDS, http://www.cbioportal.org/datasets) were added to the hypothetical database^26,27^.

### Identification and quantification of novel peptides and variant peptides by IPAW

The proteogenomics search was performed using a previously published pipeline ^4^. Briefly, all MS/MS spectra were searched by MSGFPlus^28^ in target and decoy combined database. The decoy peptide was produced by reversing protein sequences in the target database. Target and decoy matched to known tryptic peptides (from Ensembl human proteins) were discarded before FDR estimation of novel peptides. Peptide matches to known proteins were removed through BLASTP. Peptides matched to mutant peptide sequences from non-synonymous SNPs or mutations were removed. Then peptides mapping to multiple genomic loci were annotated using BLAT. The retained novel peptides were considered for further analysis. Peptides detected from 31 healthy tissues were quantified by moFF tool^10^, which extract apex MS1 intensity from raw spectral files.

### Breast cancer RNA-seq data for orthogonal evidence

We search orthogonal evidence of breast cancer peptides in breast cancer RNA-seq. A Python script search the count of reads that support novel peptides from BAM files. This script needs GFF3 format file of identified peptides and BAM files input as input files. 73 RNA-seq BAM files (63 breast cancer samples and 10 normal adjacent tissues) were downloaded from TCGA (table S8). The GFF3 files of all breast cancer datasets were merged into one. The script is available at: https://github.com/yafeng/proteogenomics_python/scam_bams.py.

### Colon cancer differential expression analysis

We used the DEP R package^29^ to impute missing values for colon cancer proteomics data (PXD002137). Before statistical analysis, missing values were imputed using random draws from a manually defined left-shifted Gaussian distribution (imputation shift =1.8, scale = 0.3). For differential expression analysis, test_diff performs a differential enrichment test based on protein-wise linear models and empirical Bayes statistics using Limma.

### Neoantigen prediction strategy

The NetCTLpan 1.1 server(http://www.cbs.dtu.dk/services/NetCTLpan/) integrates prediction of peptide MHC class I binding, proteasomal C terminal cleavage and TAP transport efficiency. The prediction parameters of NetCTLpan are set as follows: the sorting threshold ≤1%, the weight proportion of c-terminal amino acid residues was 0.225, and the weight proportion of TAP transport efficiency was 0.025.

The NetMHCpan 4.0 server (http://www.cbs.dtu.dk/services/NetMHCpan/) used artificial neural networks (ANNs) to predict the binding of peptides to any MHC molecule with a known sequence. Input peptides are digested into 9-mer peptides. Human Leukocyte Antigen (HLA) supertype representative are the allele used for prediction. The prediction parameters of NetMHCpan are set as follows: sorting threshold ≤0.5%.

### Data availability

This study is based on publicly accessible datasets listed in Table S1

## Acknowledgements

This work was supported by Science, Technology and Innovation Commission of Shenzhen Municipality (No. JCYJ20160531193931852); Science and Technology Key Project of Guangdong Province, China (No. 2019B020229002). We are thankful that the Clinical Proteomic Tumor Analysis Consortium (NCI/NIH) for sharing the data and ProteomeXchange Consortium for providing an open access proteomics data repository. We would like to thank China National GeneBank (https://db.cngb.org), Shenzhen, China for providing computing resources that support the data analysis for the paper.

## Author Contributions

Y.Z. conceived and initiated the project. Y.Z., R.X., L.M. and M.Y. contributions to the acquisition, analysis, or interpretation of data. Y.Z. and R.X have drafted and revised the manuscript together.

